# Motifizer: a tool for parsing high-throughput sequencing datasets and quantitative comparative analyses of transcription factor-binding sites

**DOI:** 10.1101/2022.04.03.486862

**Authors:** Soumen Khan, Abhik Bhattacharjee, Pranav Badhe, Chinmay Dongaonkar, Yash Gaglani

## Abstract

**Background:** Comparative analysis of Chromatin immunoprecipitation followed by massively parallel sequencing (ChIP-seq), combined with transcriptomics data (RNA-seq), can provide critical insights into gene regulatory networks controlled by transcription factors in a given biological context. While multiple programs exist, which need to be individually installed and executed to generate such information, a user-friendly tool with low system requirements which combines the various computational processes into one package, using a docker container, is lacking.

**Results:** In this study, we present a user-interactive, infrastructure-independent computational pipeline, called Motifizer, which uses open source tools for processing ChIP-Seq and RNA-seq datasets. Motifizer performs extensive analysis of raw input data, performing peak calling, de-novo motif analysis and differential gene expression analysis on ChIP and RNA-seq datasets. Additionally, Motifizer can be used for analysis and quantitative comparison of transcription factor binding sites in user defined genomic regions. Motifizer also allows easy addition and/or changes of parameters, thereby adding to the versatility of the tool.

**Conclusion:** The Motifizer tool is an easy to use tool which uses a docker container system to install and execute ChIP-seq and RNA-seq data parsing. The Analysis module of Motifizer can be employed to identify putative targets involved in gene regulatory networks. Motifizer can be accessed by a large user base and does not require programming skill by the user. Motifizer can be locally installed from the repository (https://github.com/abhikbhattacharjee/Motifizer)

## Background

The complete repertoire of interactions between a particular transcription factor (TF) with its target genes constitute a gene regulatory network (GRN). Since GRNs are critical in development and differentiation (and in disease research), identifying target genes of a TF, or the TF targeting a given gene, is important to understand the regulatory basis of biological processes. Characterization of GRN of a TF in a given biological context involves two major steps: 1) Identifying the chromatin occupancy of the TF in the genome 2) Correlation of the chromatin occupancy to the regulation of its target genes. Chromatin occupancy profile of a TF is obtained by fixing proteins onto the DNA using a chemical agent and selectively purifying the chromatin using antibodies followed by sequencing (ChIP-seq)(reviewed in Park, 2009). On the other hand, the correlation of chromatin occupancy to the regulation of a target gene by a TF comes from differential gene expression (DGE) analysis performed on transcriptomic data (RNA-seq) (reviewed in Wang, Gerstein and Snyder, 2009). Overlaying chromatin occupancy information of a TF with the regulation status of its target genes, combined with the precise location and abundance of the TF in target enhancer sequences, not only provides the gene regulatory network of the TF, but also sheds critical insights into the mechanisms of transcriptional regulation.

The analysis of ChIP-seq and RNA-seq data involves various steps performed by individual programs (Zhang *et al*., 2008; Li and Durbin, 2009; Li *et al*., 2009, 2009; Robinson, McCarthy and Smyth, 2010; Anders, Pyl and Huber, 2015; Kim *et al*., 2019). However, most of these tools require some prior knowledge of programming language or computational dependencies and requirements. While certain programs exist which provide a detailed analysis pipeline for ChIP and RNA seq datasets (Liu *et al*., 2011; Afgan *et al*., 2018; Batut *et al*., 2021), we wanted to build a software which will be fairly simple in terms of initial analysis, and therefore, more accessible to the user base with no programming expertise. In addition, while individual programs exist to calculate the occurrence and/or presence of transcription factors (Grant, Bailey and Noble, 2011), we could not find a software which could quantify and compare TF occupancy information between two or three given set of sequences. In this context, we have developed an easy to use tool which when deployed by the user using a docker container on a Linux platform, installs the various open source packages required for analysis of ChIP-seq and RNA-seq data. While the parameters used for alignment and processing of bam files are set to default, the shell scripts for such processes can be easily changed to incorporate changes as required by the user. In addition, we introduce an Analysis module which employs the FIMO software of Meme Suite to analyze and compare TF motif occupancy information in user provided genomic sequences. Taken together, we provide a versatile tool which can be employed by the larger user base for parsing of sequencing datasets and motif-based analysis of genomic sequences.

## Implementation

### Motifizer Architecture

We developed the Motifizer software with an aim to make it independent of the computational infrastructure and expertise. To maximize ease of use, the pipeline incorporates all the pre-requisite tools baked into a Docker [1] image along with custom Motifizer scripts required for data processing. The pipeline ensures modularity and provides entry points to ChIP-seq and RNA-seq processes at multiple stages. The docker image can be built at the user’s end by cloning the Dockerfile available at (https://github.com/abhikbhattacharjee/Motifizer) and deploying it using just three commands (view Readme file).

All processes carried out by Motifizer are written as a combination of shell scripts and python scripts, thereby ensuring that each process is executed successfully inside an isolated environment using docker container. To reduce the complexity of working on a CLI, a web based interactive GUI is provided based on the Flask framework. The modular design of the program allows experienced users to obtain more conclusive insights by altering the parameters/flags in shell scripts according to their needs (Supplementary information 1).

In terms of computational needs, Motifizer can be deployed on a single computing node or any platform where the underlying hardware has higher potential to obtain faster results by directly deploying the docker image. To reduce the time consumed by some of the heavy processes, parallel execution in our program using GNU Parallel (Tange, 2018) has been incorporated, which helps to reduce the processing time. The Motifizer software can be run on any engine capable of running Docker containers.

### Motifizer Functionality

The Motifizer software is divided into three main modules; a) The ChIP-seq module b) The RNA-seq module c) The Analysis module (schematic overview provided in Fig 1A, 1B, 1C). The ChIP-seq module was built using field-standard tools which involve alignment of the raw files to the reference genome using the BWA software (Li and Durbin, 2009), conversion and filtering of bam files using SAMtools (Li *et al*., 2009; Li, 2011), peak calling using the MACS2 software (Zhang *et al*., 2008) and peak annotation and de-novo motif analysis using the Homer software (Heinz *et al*., 2010). The ChIP-seq module is further divided into a) Alignment module b) Peak calling and annotation module. The Alignment module requires single ended fastq files (test and input) and the reference genome fasta as input, and provides the bam files, peak file, annotation of the peak file and de-novo motif enrichment analysis as output (Fig 1A). The Peak calling and Annotation module, on the other hand, skips the alignment step and requires bam files as input. While all modules have been developed using default parameters, the shell scripts of the executable files can be easily edited to change and/or add additional parameters (Supplementary information 1).

**Figure 1.**
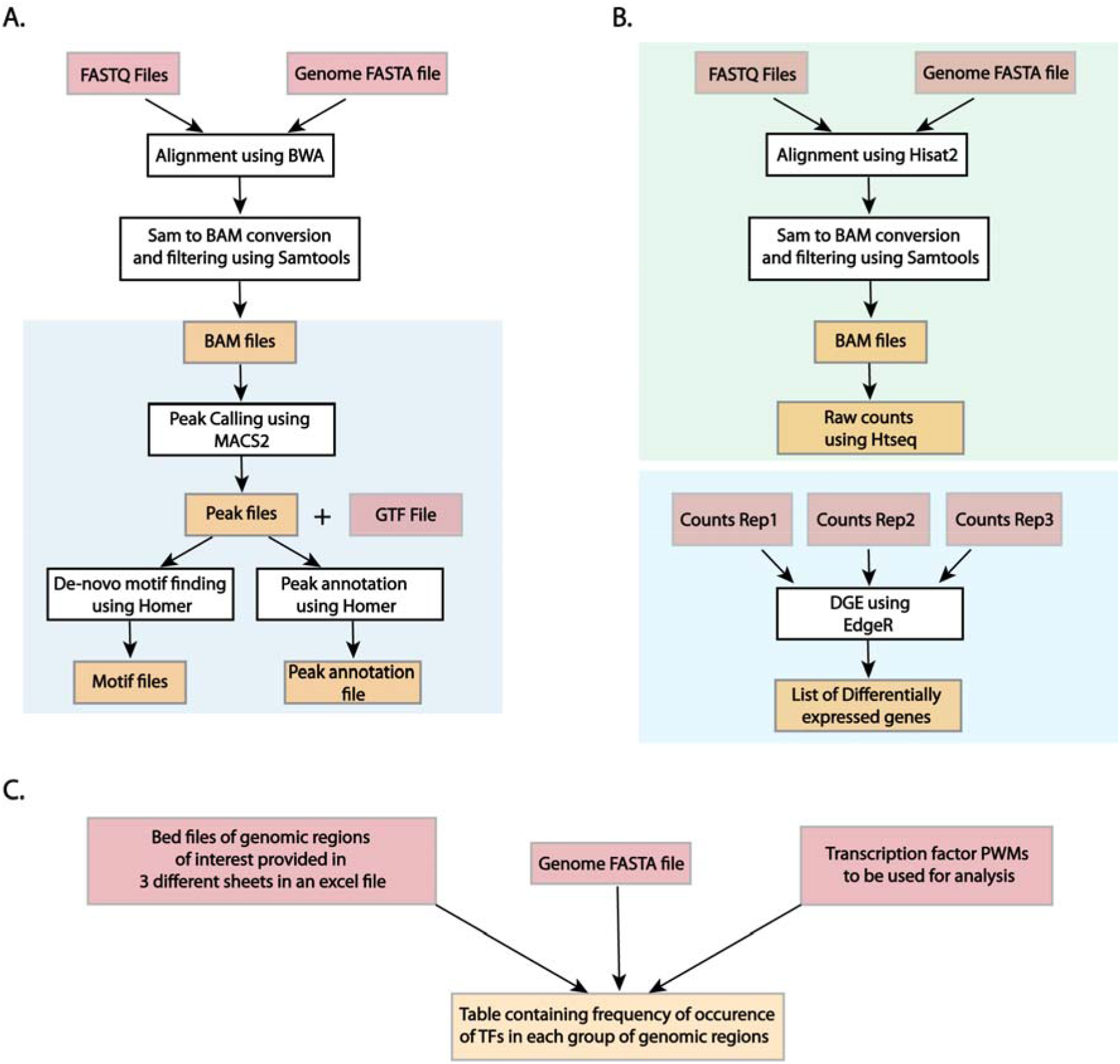
Modules for ChIP-seq, RNA-seq and Analysis provided by Motifizer. **A**. The ChIP-seq module of Motifizer requires three input files (highlighted in pink) from the user; the fastq sequences, the genome fasta file and the GTF file (annotation file) and provides four output files (highlighted in yellow); the bam files, the peak output files, the de-novo motif output and the peak annotation file. Motifizer also provides an option for peak calling and annotation by taking bam files as input (portion highlighted by the blue box) **B**. The RNA-seq module operates in two different parts. The first part (highlighted in green box) requires fastq files and the genome fasta files as input (highlighted in pink) and provides raw counts files as output ((highlighted in yellow). The second part (highlighted in blue box) requires count files as input (three biological replicates for control and test) and provides a list of differentially expressed genes as output. **C**. The Analysis module requires genomic coordinates (in bed format) and genome fasta file and Transcription factor PWMs as input and provides a table calculating the frequency of each TF in each group of enhancers provided as input.

The RNA-seq module also utilizes field standard software and performs read alignment using the Hisat2 software (Kim *et al*., 2019), filters out duplicate reads using SAMtools, performs read counting using the HTSeq software (Anders, Pyl and Huber, 2015), and employs the edgeR package (Robinson, McCarthy and Smyth, 2010) to determine differentially expressed genes. The RNA seq module is further divided into a) Alignment b) edgeR analysis. The Alignment module requires fastq files (either single or paired end) and the reference genome fasta file as inputs and provides read count files as output (Fig 1B). The edgeR module requires count files as input and provides the list of differentially expressed genes as output (Fig 1B). Similar to the ChIP-seq module, the RNA-seq module can be easily edited in a text editor before execution to change parameters (Supplementary Information 1).

One of the novel aspects of Motifizer is an Analysis module which can be used to quantify and compare the frequency of transcription factor motifs in user provided genomic sequences. The Analysis module employs the FIMO software (Grant, Bailey and Noble, 2011) to scan and count occurrences of transcription factor PWMs in genomic sequences and provides the number and frequency of occurrence as output. The Analysis module requires genomic coordinates (in bed format) in an excel format (template provided) and can be used to compare up to three different categories of genomic sequences (provided in three different sheets) (Fig 1C).

## Results

### 1. Analysis of ChIP-seq datasets using Motifizer

We ran the Alignment module on ChIP-seq data generated for Hth (Loker, Sanner and Mann, 2021) in *Drosophila melanogaster* haltere imaginal discs and for dFOXO (Birnbaum *et al*., 2019) in *Drosophila melanogaster* embryos. The total process, starting from alignment to de-novo motif analysis, was completed in less than four hours for each of the datasets. De-novo motif analysis of Hth ChIP-seq data is consistent with the finding that binding motif for Hth (Fig 2A) is enriched in ChIP pulled down sequence. For dFOXO ChIP-seq dataset, we report enrichment of reported motifs, albeit with differences in observed p-values (Fig 2B). Since all analyses were performed using default parameters, we encourage users to modify the same according to their own requirements to replicate their results.

**Fig2:**
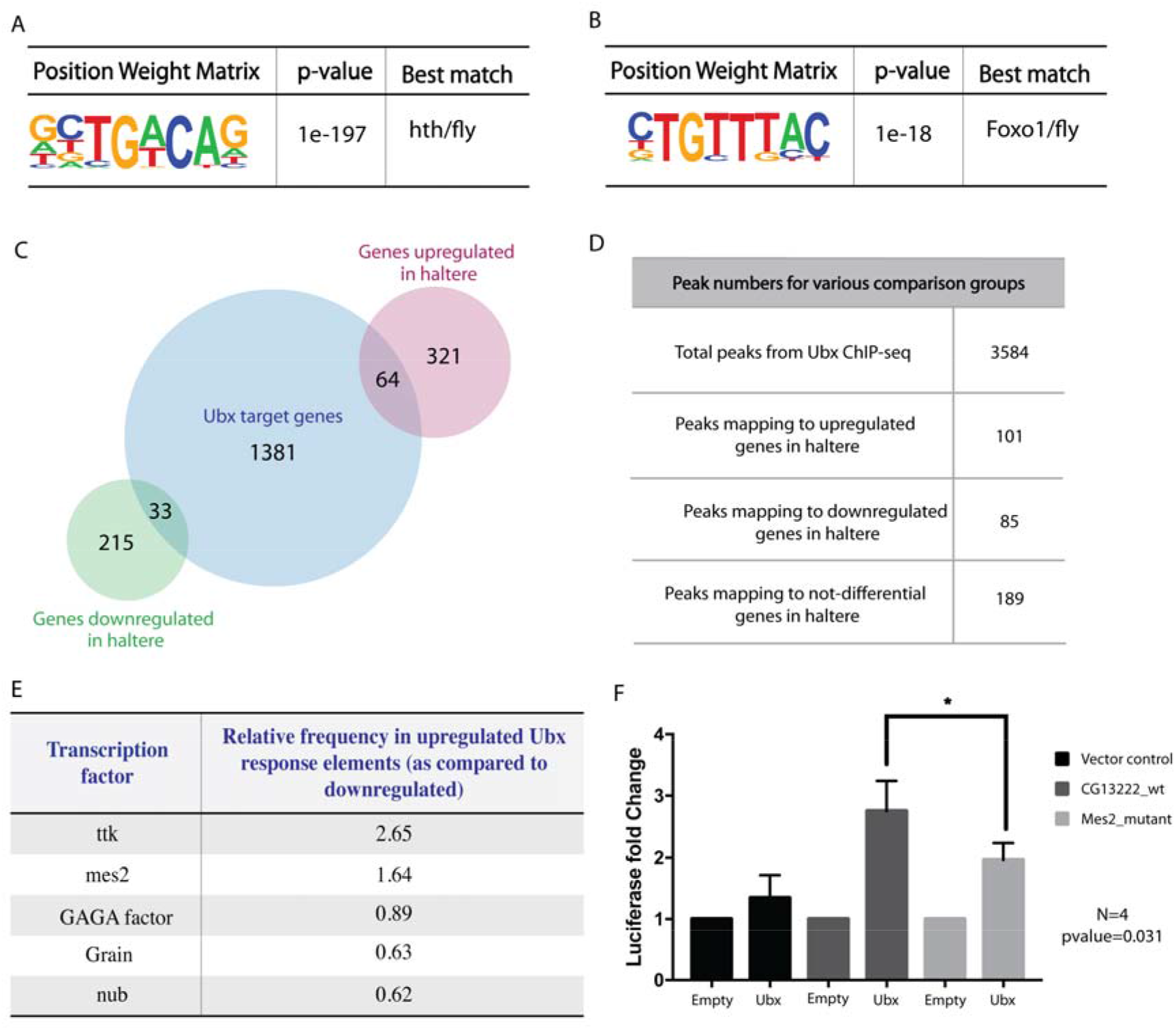
Motifizer identifies critical TFs involved in gene regulatory pathways during haltere development in *Drosophila melanogaster*. **A-B**. Analysis of Hth ChIP-seq data from *Drosophila* halteres using the ChIP-seq module of Motifizer reveals enrichment of a motif predicted to be bound by the Hth protein. Similar observations are made for dFOXO ChIP-seq data generated in Drosophila embryos **C-D**. Comparison of Ubx ChIP-seq data generated in *Drosophila* halteres (using the ChIP-seq module) with differentially expressed genes between wing and haltere imaginal discs (using RNA-seq module) suggest that 64 genes are upregulated by Ubx in the haltere, whereas 33 genes are downregulated by Ubx in the haltere (C). The 64 upregulated genes had 101 Ubx peaks associated with them whereas the 33 downregulated genes had 85 peaks associated. 189 peaks were annotated to Ubx targets which were neither upregulated nor downregulated. **E**. Comparison of TF occupancy across upregulated, downregulated and not-differential enhancers (using the Analysis module). Motifs predicted to be bound by ttk and mes2 are enriched in upregulated enhancers as compared to downregulated enhancers. **F**. The *CG13222* enhancer drives Luciferase activity in a S2 cell culture system in presence of Ubx. Significant loss of reporter activity is observed in the mutant enhancer (Mes2_mutant) where the Mes2 motif is mutated. For statistical analysis, t-test was performed (two-tailed).

### 2. Identification of transcription factor signatures in the Hox gene, Ultrabithorax, mediated haltere specification in Drosophila melanogaster

The Hox gene, Ultrabithorax, specifies the development of the halteres, a modified wing structure, in the third thoracic segment (Lewis, 1978) in *Drosophila melanogaster*. Ubx displays very high specificity of function in-vivo and is known to upregulate and downregulate a large number of genes in order to specify the haltere fate (Weatherbee *et al*., 1998). However, in-vitro, Ubx displays poor specificity and affinity of binding to a recognition sequence containing a core TAAT (Ekker *et al*., 1991; Berger *et al*., 2008; Noyes *et al*., 2008). Thus, it is hypothesized that Ubx interacts with different groups of co-transcription factors to attain specificity of function.

In an attempt to identify such co-transcription factors involved in Ubx mediated modulation of gene regulatory pathways, we developed a methodology as described in Fig 2A. We have previously generated ChIP-seq data for Ubx in third instar haltere imaginal discs. We analyzed it using the ChIP-seq module of Motifizer. Next, we combined the three biological replicates to identify high confidence peaks and annotated them using the Homer software (detailed in Supplementary Materials and Methods). We have also previously generated RNA-seq data for wing and haltere imaginal discs. We analyzed it using the RNA seq module and identified differentially expressed genes between the wing and haltere imaginal discs. We compared the annotated peaks file (obtained from ChIP-seq analysis) to differentially expressed genes and identified 64 genes which were Ubx targets and upregulated in the haltere and 33 genes that are downregulated in the haltere (Fig 2C). These further correspond to 101 and 85 genomic regions bound by Ubx which annotated to genes that are upregulated or downregulated in the haltere, respectively (Fig 2D). In addition, we also identified 189 genomic regions which annotated to genes that are not-differentially expressed (0.95<FC<1.05) between the wing and halteres. We ran the analysis module using all known *Drosophila* transcription factors (JASPAR database) and quantitated the frequency of occurrence of each TF (Fig 2E). We find that known cofactors of Ubx like MAD and GAGA factor are enriched in enhancers of downregulated targets. Interestingly, we found that a motif predicted to be bound by the Mes2 transcription factor is enriched specifically in Ubx bound regions that annotate to genes that are upregulated in the haltere. In this context, we analyzed the enhancer region of a gene, CG13222, which is upregulated by Ubx and identified a Mes2 binding site in close proximity to the Ubx binding site. In a Luciferase based reporter assay system, we observe that while the wild type CG13222 shows a ∼3-fold increase in reporter expression on Ubx induction, a mutant CG13222 enhancer with a mutated Mes2 motif, shows a significantly less activation of the enhancer (Fig 2F). This suggests the possible role of the Mes2 enhancer in the Ubx mediated activation of the CG13222 enhancer, thereby validating the role of the analysis module in identifying putative targets essential for regulation of gene regulatory networks.

## Conclusion

We provide an easy to use software requiring minimal computational memory which can be used by the larger user base without any prior knowledge of any programming language. Motifizer presents a quick and reliable solution for parsing ChIP-seq and RNA-seq datasets and can be effectively used to provide motif-based analysis of transcription factors involved in a given gene regulatory network.

## Availability and requirements

- Project name: Motifizer
- Project home page: https://github.com/abhikbhattacharjee/Motifizer
- Operating system: Linux
- Any restrictions to use by non-academics: None
- Dependencies: Docker container

## Acknowledgments

We thank LS Shashidhara for providing critical inputs during project conception as well as facilities for library preparation, sequencing and analysis platforms. We thank Leelavati Narlikar for her guidance, and Guillaume Giraud and Samir Merabet for critical inputs. This work was supported primarily by a JC Bose Fellowship to LS Shashidhara and a Council of Scientific & Industrial Research (CSIR) Fellowship to SK.

## Author contributions

SK carried out the ChIP-seq, RNA-seq, Luciferase assay and designed the methodology.

AB and PB wrote the code, built the GUI and docker container and performed test runs.

CD and YG contributed to writing the code and performed test runs.

SK conceived the project and wrote the MS.

