## Supplementary Information for "Motifizer: a tool for parsing high-throughput sequencing datasets and quantitative comparative analyses of transcription factor-binding sites"

**Supplementary Information, Figures and Materials and Methods**

**Supplementary Information 1: Changing parameters in .sh files**

The modularity incorporated into the pipeline allows the user to change configurations specific to his requirements. The commands for following modules can be moulded by an experienced user by modifying the open-source code available at <https://github.com/abhikBhattacharjee/Motifizer>

The following section highlights the files and part of the code written in such files that can be modified by the user. The code highlighted in green can be edited but not the grey highlights.

**1. ChIP-Seq Module**

**File names:** ChIP_seq.sh & chip_annotation.sh

**Part of the code that can be modified:**

**a.** For Peak Calling

$macs2 callpeak -t bwa_test_filtered.bam

-c bwa_input_filtered.bam

--outdir macs2

--format AUTO

--gsize dm

--pvalue 0.05

--call-summits

Users can provide the specific format of the input files (default has been set to AUTO). The genome size, p-value and other options can be changed or added depending on the requirement of the user. More details about the MACS2 parameters can be found here: [https://manpages.ubuntu.com/manpages/xenial/man1/macs2_callpeak.1.html[1](https://manpages.ubuntu.com/manpages/xenial/man1/macs2_callpeak.1.html%5b1)]

b. For peak annotation

$ annotatePeaks.pl macs2/NA_summits.bed

none

-gtf $4

-noblanks

> homer_data

- Additional parameters for annotation can be added. Further details available here: <http://homer.ucsd.edu/homer/ngs/quantification.html>[2]

b. For denovo motif analysis

$ findMotifsGenome.pl macs2/NA_summits.bed

$1

output_motif

-mask

-size 200m

-mset insects

- Additional parameters for denovo motif analysis can be added. Further details available here: [http://homer.ucsd.edu/homer/ngs/peakMotifs.html](http://homer.ucsd.edu/homer/ngs/peakMotifs.html%5b2)[2]

**2. RNA-Seq Module:**

**File names: rna_hisat_se.sh**, **rna_hisat_pe.sh &**

**Part of the code that can be modified:**

a. For read counting:

$ htseq-count

--format=bam

--stranded=no

--type=exon

hisat2_filtered.bam $4 > $5

- Additional parameters for can be added. Further details available here: [https://htseq.readthedocs.io/en/release_0.11.1/count.html[6](https://htseq.readthedocs.io/en/release_0.11.1/count.html%5b6)]

**Supplementary Information 2: Guidelines to make the excel file for analysis**

To prepare the file for implementation of the analysis module, a template excel file has been provided. This file contains three sheets labelled as ‘Enhancer_Group1’, ‘Enhancer_Group2’, and ‘Enhancer_Group3’. The excel sheet prepared and used for the results shown in this study has also been provided as Supplementary data.

a. For calculating the frequency of binding of user provided transcription factors, genomic coordinates of the sequences need to be provided by filling any one of the sheets. The remaining two can remain empty.

b. For calculating and comparison of binding of user provided transcription factors between different enhancer groups, sequences need to be provided by filling two or three sheets (depending on the number of enhancer groups).

**Supplementary Materials and Methods**

**1. Comparative analyses of putative Ubx response elements:** Genes that are differentially expressed between the wing and haltere were identified using RNA sequencing of third instar wing and haltere imaginal discs. A fold difference (FD) cut-off of 1.5 fold was applied to identify upregulated or downregulated genes in the haltere. Genes having a fold difference ranging between 0.95-1.05 between the wing and haltere imaginal discs were considered as not-differentially expressed. Ubx targets obtained from ChIP-seq was compared to RNA-seq data and frequency of the TAAAT motif calculated using an analysis module (unpublished data).

**2. Molecular Cloning:** Site directed mutagenesis approach was used to mutate the Mes2 binding motif in the CG13222 enhancer using the following primer: 5’-TTGTCCTTCTGTCAATGTTTTGGAAAAAACCAATAAAGCTG-3’ and cloned between Kpn1 and Nhe1 restriction sites, upstream of a modified pGL3 vector containing a 5X Dorsal binding site. Metallothionein inducible pRMHa3 vectors containing Ubx were previously generated in the lab and empty pRMHa3 vector was generated by excising the cloned Ubx sequence. All constructs were sequence verified before use in functional assays.

**3. Luciferase assays:** S2 cells were checked for contamination before performing Luciferase assays. Cells were plated onto 24 well plates at a density of 3*10^5 cells per well 6 hours prior to transfection. For every construct, either wild type or mutant, two sets of experiments were designed; one well was co-transfected with the enhancer construct in pGL3 vector and the empty pRMHa3 vector whereas the other well was co-transfected with the enhancer construct and pRMHa3 vector containing Ubx. Renilla luciferase was used as an internal control and co-transfected in all experiments.  Transfection was carried out using the Effectene transfection reagent and all experiments carried out in 3 technical replicates and at least 3 biological replicates. 48 hours post transfection, sterile CuSO4 solution was used to induce expression of Ubx at a final concentration of 500um and incubated for 24 hours. Cells were harvested, pelleted down (1000rpm for 4mins) and 100ul of 1X Passive Lysis buffer added and vortexed to dissolve the cell pellet. Cells were incubated for 15mins at room temperature and further spun at 10,000rpm for 90secs to collect the supernatant. The luminescence was measured with the Dual-glo Luciferase assay kit (Promega) on Ensight Plate reader (Perkin Elmer). All readings were normalized to Renilla luminescence and datasets compared using the GraphPad Prism software.
